# Putting RFMix and ADMIXTURE to the test in a complex admixed population

**DOI:** 10.1101/671727

**Authors:** Caitlin Uren, Eileen G. Hoal, Marlo Möller

**Author notes:** Correspondence should be addressed to: Dr. Caitlin Uren, Room 4036, 4^th^ Floor Education Building, Francie van Zijl Drive, DST-NRF Centre of Excellence for Biomedical Tuberculosis Research, South African Medical Research Council Centre for Tuberculosis Research, Division of Molecular Biology and Human Genetics, Faculty of Medicine and Health Sciences, Stellenbosch University, Cape Town, 8000, South Africa.

## Abstract

Global and local ancestry inference in admixed human populations can be performed using computational tools implementing distinct algorithms, such as RFMix and ADMIXTURE. The accuracy of these tools has been tested largely on populations with relatively straightforward admixture histories but little is known about how well they perform in more complex admixture scenarios. Using simulations, we show that RFMix outperforms ADMIXTURE in determining global ancestry proportions in a complex 5-way admixed population. In addition, RFMix correctly assigns local ancestry with an accuracy of 89%. The increase in reported local ancestry inference accuracy in this population (as compared to previous studies) can largely be attributed to the recent availability of large-scale genotyping data for more representative reference populations. The ability of RFMix to determine global and local ancestry to a high degree of accuracy, allows for more reliable population structure analysis, scans for natural selection, admixture mapping and case-control association studies. This study highlights the utility of the extension of computational tools to become more relevant to genetically structured populations, as seen with RFMix. This is particularly noteworthy as modern-day societies are becoming increasingly genetically complex and some genetic tools are therefore less appropriate. We therefore suggest that RFMix be used for both global and local ancestry estimation in complex admixture scenarios.

## Introduction

Admixture, the exchange of genetic material between distinct populations, is a hallmark of modern society - it can occur between closely or distantly related populations (both genetically and geographically) (1000 Genomes Project Consortium *et al.* 2012). This exchange of genetic material leads to population structure; the pattern, timing and extent has been investigated in detail in a number of populations (1000 Genomes Project Consortium *et al.* 2012; Gurdasani *et al.* 2015; Uren *et al.* 2016). Such studies on southern African populations are particularly noteworthy as this area is postulated to be the geographical origin of modern humans and therefore investigating population structure in modern southern African populations may reveal more about the area’s rich history (Henn *et al.* 2011).

Correctly and efficiently determining ancestral proportions in an admixed population is possible by using computational and statistical algorithms that adapt to a variety of demographic scenarios (Alexander *et al.* 2009; Maples *et al.* 2013; Brown and Pasaniuc 2014). Furthermore, the ability to determine the ancestral origin of a particular chromosomal region in an admixed individual has enabled the mapping of the origins of genetic risk factors in complex disease i.e. admixture mapping (Freedman *et al.* 2006; Cheng *et al.* 2009; Daya *et al.* 2014). The majority of the computational and statistical tools used for global and local ancestry were however tested on and tailored to 2- to 3-way admixed populations. The extension to a complex 5-way admixed population and the evaluation of the resulting accuracy as we present here, has rarely been done. (Daya *et al.* 2014; Uren *et al.* 2016).

A South African population with unique genetic ancestry and 5-way admixuture (the South African Coloured (SAC) population as termed in the South African census) received ancestral contributions from Bantu-speaking African (~30%), KhoeSan (~30%), European (~20%), East Asian (~10%) and South East Asian populations (~10%) (de Wit *et al.* 2010; Chimusa *et al.* 2013; Uren *et al.* 2016). The admixture began approximately 15 generations ago and followed a continuous migration model (Uren *et al.* 2016). This number, mode and timing of admixture events is unique and creates a highly complex population. Although we are able to describe the demographic model for most populations, there are some gaps in knowledge. This may include not knowing which populations or specific geographical locations are the best proxies for the true ancestral populations. Therefore, any studies investigating an association between genetics and disease risk needs to be able to correctly account for population or even individual admixture proportions within the limits of the availability of current genetic data.

The first step in a study design aimed at finding a link between ancestry and disease (such as genome-wide association studies and admixture mapping) is to understand the ancestral composition of the study population. Ancestral origins and contributions to the 5-way admixed South African population have been estimated but there have been very few studies that have investigated the accuracy of the results generated by the computational algorithm used (de Wit *et al.* 2010; Chimusa *et al.* 2013; Petersen *et al.* 2013; Daya *et al.* 2013; Uren *et al.* 2016). Here we have set out to test the accuracy of global and local ancestry inference in one of the most complex admixed populations world-wide, using newly available dense genotyping data. A simulated 5-way admixed population is generated and global and local ancestry estimates are compared to the true values to determine the accuracy of the computational algorithm.

## Methods

### Data merging and filtering

KhoeSan genotype data from Martin and colleagues (Martin *et al.* 2017) was merged with the PAGEII dataset (Wojcik *et al.* 2018). In order to increase the number of European and South East Asian reference samples in the dataset, the data was merged with Gujarati Indian and European genetic data from the 1000 Genomes Project (1000 Genomes Project Consortium (2010) 2010).

Preliminary data filtering included a filter for minor allele frequency (0.003), missingness per genotype (max 0.05) and missingness per individual (max 0.01). A total of ~776k SNPs passed these filters and formed the initial merged dataset. Further data filtering is described in the appropriate sections below. Data was phased using SHAPEIT2 utilizing a recombination map averaged across European and African populations (The International HapMap Consortium 2007; O’Connell *et al.* 2014). A summary of the populations in the final dataset can be seen in Table 1.

**Table 1:**
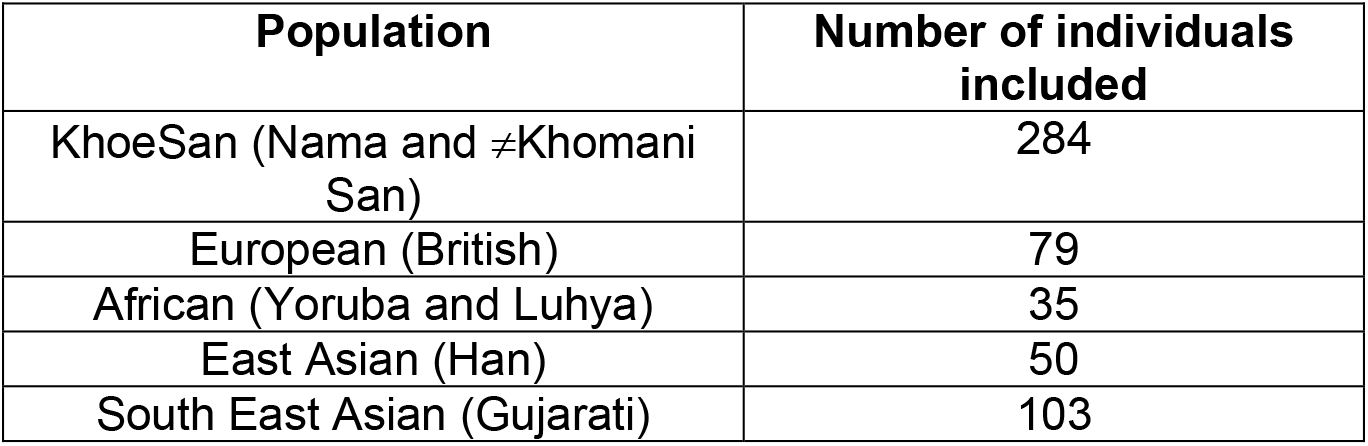
Population characteristics of the final merged dataset.

### Simulations

The computational workflow is summarized in Figure 1. A subset of reference individuals from the final merged dataset described in Table 1 was used to generate a simulated dataset using admix-simu (Williams 2016). A demographic model consisting of the ancestry proportions described above and a continuous migration model starting at 15 generations ago (Uren et al. (2016)), was used to generate a simulated 5-way admixed population (Uren *et al.* 2016). This simulation results in a heterogenous population, reminiscent of a real-world SAC population. The simulation does not take post-admixture selection into account since it is highly unlikely that 350 years would result in distinct selection signals, rather, the inherent selection signals in the source populations will be transferred in a random manner to the simulated admixed population. Genotype as well as local ancestry calls were generated for this simulated dataset from real reference haplotypes, thus capturing the complexity of this heterogenous 5-way admixed South African population.

**Figure 1:**
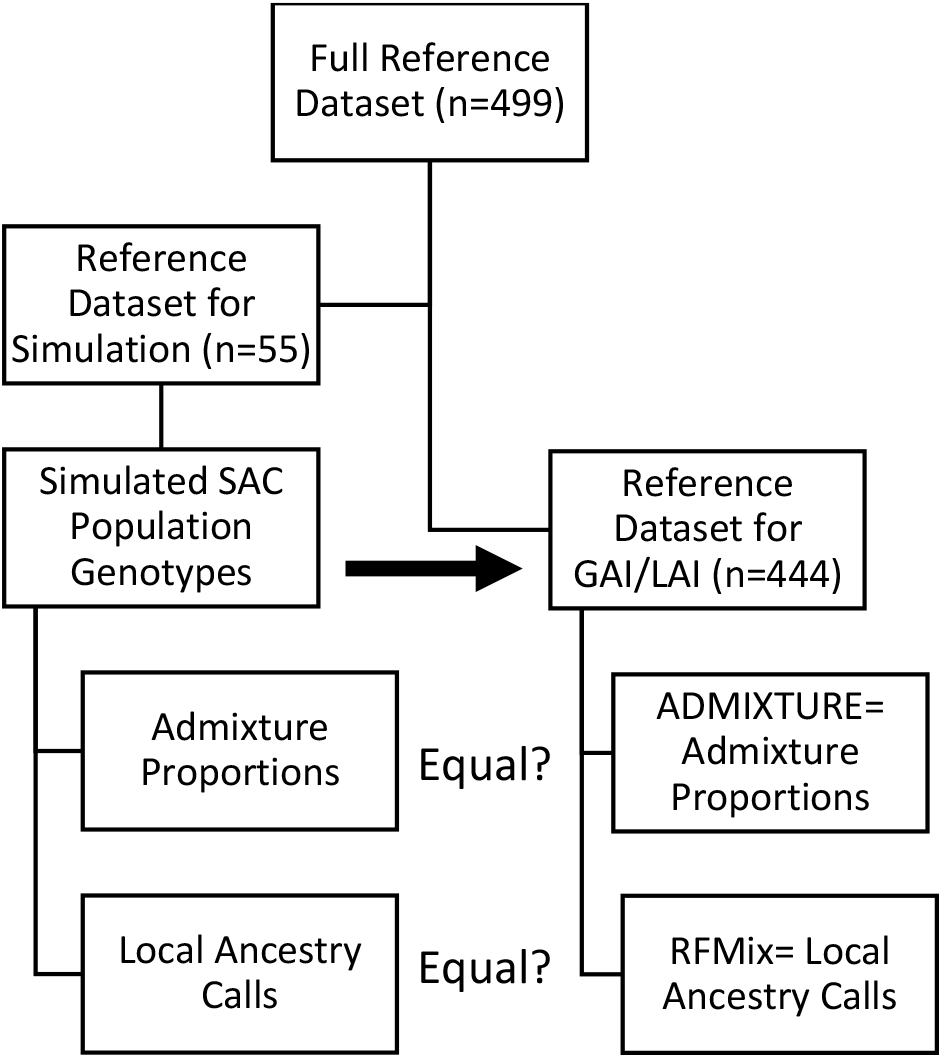
Computational workflow. The number (n) of individuals included in each dataset, over all ancestral populations. For details, please see the methods section.

### Software choices

Although there are a number of software programs that are able to estimate global ancestry, ADMIXTURE is the most utilized. Reasons for this include the ability to include related individuals in one run and to generate accurate admixture proportions using relatively low-density SNP-array data (Alexander *et al.* 2009). The other widely used global ancestry algorithm, STRUCTURE has been shown to overestimate admixture proportions in complex populations (Cheng *et al.* 2017).

RFMix was chosen as the local ancestry inference algorithm of choice as it allows for parameter optimization given the number of ancestral populations, has the inherent ability to calculate local and global ancestry simultaneously, allows for array-based input data as well as whole genome sequencing data, and has a proven track record with admixed populations (Maples *et al.* 2013; Padhukasahasram 2014). Furthermore, during a preliminary study by Daya and colleagues, RFMix was shown to be the most accurate tool for local ancestry estimation in this 5-way admixed South African population (Daya *et al.* 2014).

### GAI accuracy

Reference individuals not included in the dataset used for the simulation, were allocated to the dataset used for global and local ancestry inference. Global ancestry proportions were determined by ADMIXTURE (Alexander *et al.* 2009) and RFMix (Maples *et al.* 2013).

The ADMIXTURE analysis was performed in an unsupervised manner after filtering the dataset for linkage disequilibrium as per recommendations in the ADMIXTURE manual (50kb window size, step size of 10kb and R^2^ threshold of 0.1). Relatedness in the reference dataset was assessed using king (Manichaikul *et al.* 2010) and all second degree relatives were removed prior to admixture analysis.

RFMix was run using default parameters, a time since admixture of 15 generations (in line with the simulation) as well as 3 expectation-maximization (EM) iterations (further EM iterations were not shown to increase accuracy (Maples *et al.* 2013)). The correlation of the two methods by means of the Root Mean Squared Error was performed in R.

### LAI accuracy

Local ancestry calls were generated by RFMix using the same parameters as described in the previous section.

The ability to correctly assign local ancestry was calculated in two ways. The first determined the global accuracy i.e. how often the computational tool assigned the correct ancestry (as per the simulations) and the second looked at this accuracy per ancestral population (Atkinson 2018). These accuracy estimators were then averaged over all individuals in the simulated 5-way admixed dataset.

### Data Availability

No new genetic data was generated for this study however all reference data supporting the findings of this study are available via the original publication.

## Results and discussion

The aim of this study was to determine the accuracy of global and local ancestry inference. In order to do this, a highly complex 5-way admixed population was simulated. The local and global ancestry estimates were then compared to the true simulated data.

### Global Ancestry Inference Accuracy

The genetic diversity inherent in an admixed South African population was simulated using 5 reference populations (see Methods). The average ancestry proportions across these individuals were in line with what is seen in the real-world (Table 2) (Uren *et al.* 2016). The simulations provided the basis with which the global ancestry proportions as calculated by ADMIXTURE (Alexander *et al.* 2009) and RFMix (Maples *et al.* 2013) could be compared.

**Table 2:** Average admixture proportions.

|  | Previously Reported (Uren *et al.* 2016) (%) | Simulation (%) | ADMIXTURE (%) | RFMix (%) |
| --- | --- | --- | --- | --- |
| Bantu-speaking African | 32 | 27 | 33 | 26 |
| KhoeSan | 30 | 33 | 25 | 33 |
| European | 19 | 22 | 26 | 23 |
| East Asian | 7 | 6 | 7 | 6 |
| South East Asian | 12 | 12 | 9 | 12 |

Unsupervised admixture analysis of the simulated dataset by ADMIXTURE and RFMix confirmed that the simulated 5-way admixed population is highly heterogenous. Average ancestral proportions for both computational tools are given in Table 2. The comparisons across the 5 ancestries for each simulated individual are also depicted in Figure 2. Root Mean Squared Errors (RMSE) (RFMix vs Simulation and ADMIXTURE vs simulation) were calculated for each ancestry. As per the RMSE’s, RFMix outperforms ADMIXTURE in correctly estimating admixture proportions in the 5-way admixed population, with the exception of East Asian ancestry where the accuracy is equal. ADMIXTURE over-estimates the Bantu-speaking African contribution and under-estimates the KhoeSan ancestral proportions. ADMIXTURE also overestimates European ancestry and underestimates South East Asian ancestry. This is most likely due to inherent European ancestry present in South East Asian populations and similarly, Bantu-speaking ancestry in the KhoeSan reference population. It is likely that if more homogenous reference populations were chosen, this trend would be negated but, as previously mentioned, most modern day populations are admixed and therefore computational tools should be able to account for this within the algorithms.

**Figure 2:**
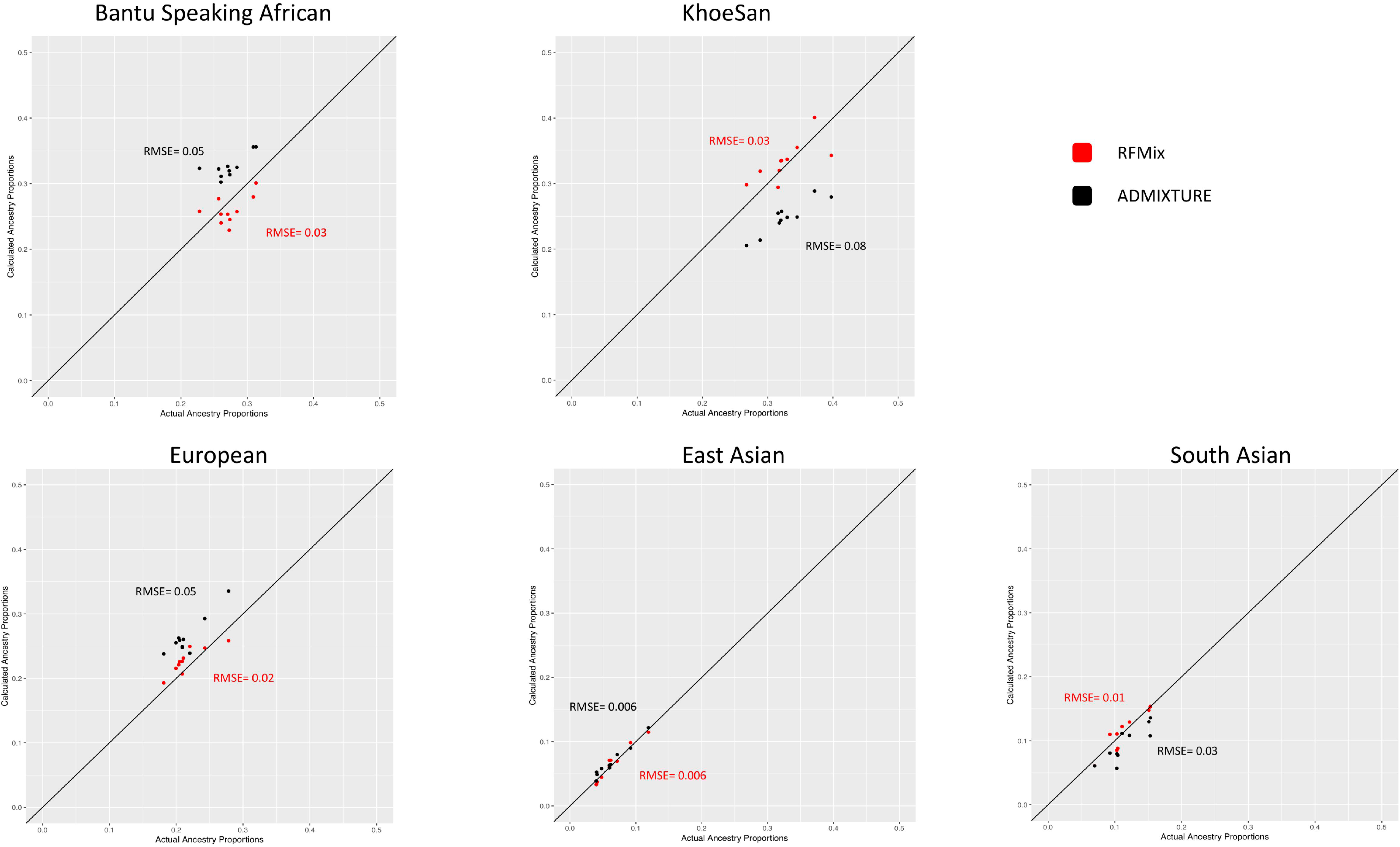
Comparison between observed global ancestry proportions and “true” proportions showing RFMix performs more accurately than ADMIXTURE in ancestry determination. Admixture proportions calculated by ADMIXTURE are in black and RFMix in red. Root Mean Square Errors for every comparison are shown.

In addition, we hypothesize that the discrepancy in admixture proportions between RFMix and ADMIXTURE can also be attributed to the increase in prior information given to RFMix in order to determine admixture proportions i.e. phase and recombination rate.

### Local Ancestry Inference Accuracy

Beyond global ancestry proportions, the simulation of a 5-way admixed population resulted in known local ancestry calls, to which calls by a computational tool can be compared. The ancestral origin of each parental chromosomal region was determined using RFMix. RFMix has been shown to outperform other computational tools in the estimation of local ancestry in complex admixture scenarios (Daya *et al.* 2014). The local ancestry calls by RFMix were compared to the “true” simulated ancestral origin of each region (Figure 3). The overall local ancestry inference accuracy across all individuals and ancestries is ~89%; 88% accurate in calling Bantu-speaking African ancestry, 87% calling KhoeSan ancestry, 95% calling European ancestry, 86% calling East Asian ancestry and 85% calling South East Asian ancestry. The statistical significance of RFMix’s ability to call a specific ancestry over another was assessed. RFMix is able to call European ancestry more precisely than any of the African or Asian ancestries and it was able to call KhoeSan ancestry more accurately than East Asian ancestry (Figure 3). This is consistent with what was previously found and confirms that these algorithms are not tailored to African populations (Maples *et al.* 2013).

**Figure 3:**
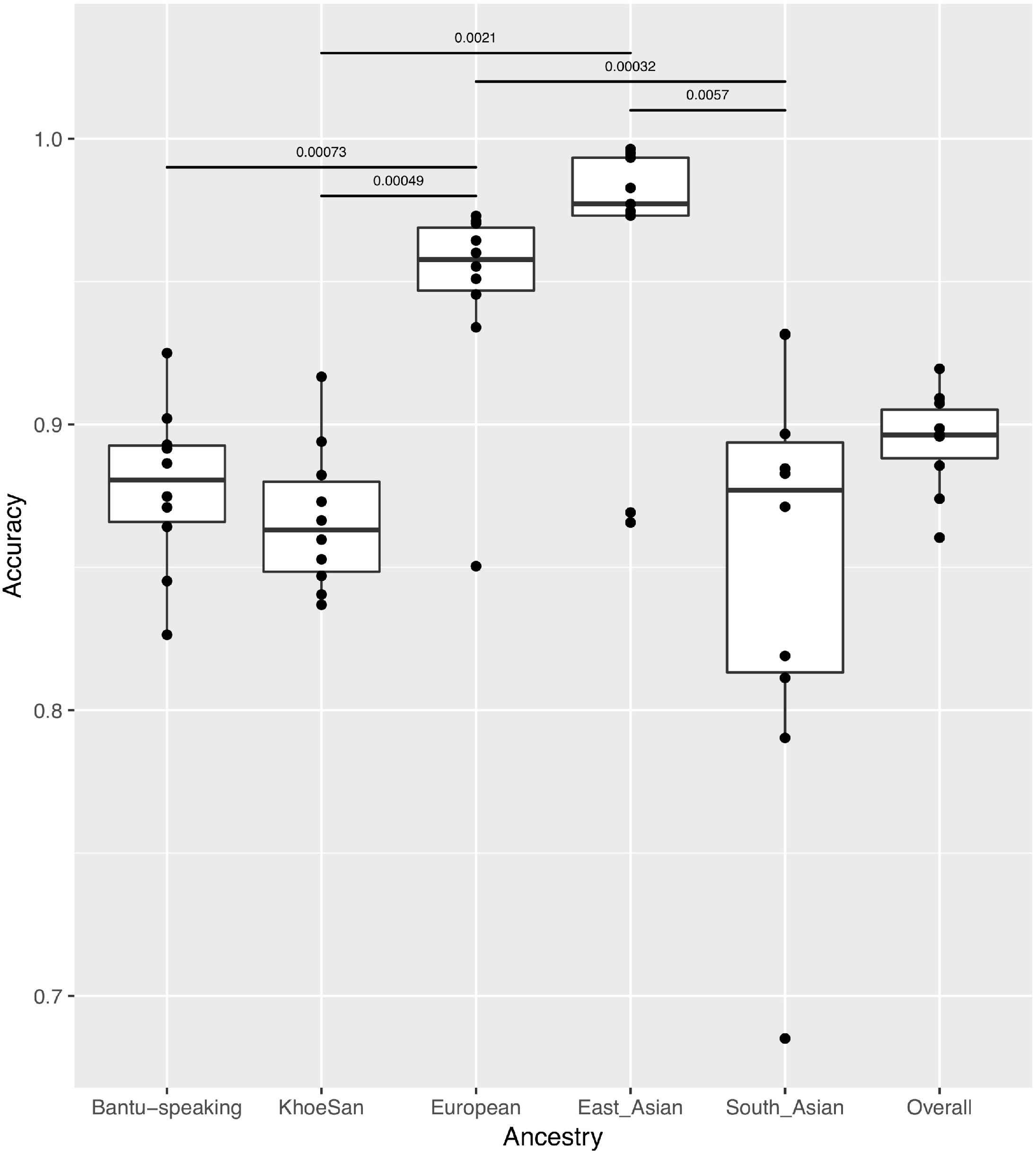
Boxplot showing the accuracy with which RFMix assigns an ancestral origin to a genetic region, stratified by reference population. The median (bold horizontal line) and the upper and lower quartiles are shown. Data faliing outside this range are plotted as outliers. The differences in accuracies across ancestries were assessed using a Wilcoxon non-parametric test. All statistically significant p values (< 0.01) are shown.

The local ancestry inference accuracy estimates presented here are substantially higher than previously obtained for this 5-way admixed South African population (Daya *et al.* 2014). This increased accuracy can be attributed largely to the recent availability of large-scale genotyping array data from the KhoeSan population which is used as a reference for this admixed population. This, in addition to the overall higher SNP density, increased the accuracy from ~70% (as previously reported (Daya *et al.* 2014)) to ~89%. As new datasets become available and the overlap between datasets increases, we envisage this accuracy increasing even further.

## Conclusion

In conclusion, the findings presented here detail the accuracy of global and local ancestry inference of one of the most complex populations worldwide, which puts ADMIXTURE and RFMix to the ultimate test. Due to the accuracy and versatility of RFMix in determining global and local ancestry in a single program, it should be the algorithm of choice to characterize more complex admixture scenarios. The inclusion of accurate admixture proportions as a covariate in association studies is vital, and it is our opinion that researchers studying complex admixed populations should use RFMix for this purpose.

Furthermore, we demonstrate that computational tools *are* able to decipher the complex African genetic history with a high degree of accuracy, but there is still some room for improvement regarding the tailoring of computational tools to handle diverse, admixed reference and target populations under study.

As populations become increasingly mobile, the probability of admixture is greater and the extension of these and future computational tools to more genetically complex populations across the world is vital and, as we have demonstrated, is possible. The conclusions of this study will be increasingly relevant and generalizable.

## Acknowledgements

We thank Dr. Brenna Henn and Dr. Elizabeth Atkinson for their assistance and support of this research. We thank the study participants of the projects cited here - without their contribution, this research would not be possible.

## Author Contributions

CU designed the study, wrote the first draft of the manuscript and performed the computational analyses. MM and EH helped to develop the research and edited the manuscript.

## Funding

This research was funded (partially or fully) by the South African government through the South African Medical Research Council and the National Research Foundation. CU was supported by a fellowship from the Claude Leon Foundation.

## Conflict of Interest

The authors declare that they have no conflict of interest.

## Ethical Approval and Informed Consent

All procedures performed in studies involving human participants were in accordance with the ethical standards of the institutional and/or national research committee and with the 1964 Helsinki declration and its later amendments or comparable ethical standards. Informed consent was obtained from all individual participants included in the study. This study was approved by the Stellenbosch University Health Research Ethics Committee (Reference #N11/07/210).

